# Debunking the “junk”: Unraveling the role of lncRNA–miRNA–mRNA networks in fetal hemoglobin regulation

**DOI:** 10.1101/2021.10.13.464339

**Authors:** Motiur Rahaman, Chiranjib Bhowmick, Jaikrishna Komanapalli, Mandrita Mukherjee, Prasanna Kumar Byram, Praphulla Chandra Shukla, Tuphan Kanti Dolai, Nishant Chakravorty

**Affiliations:** School of Medical Science and Technology, IIT Kharagpur, Kharagpur, Paschim Medinipur, West Bengal – 721302, India; Department of Biotechnology, IIT Kharagpur, Kharagpur, Paschim Medinipur, West Bengal – 721302, India; Department of Hematology, Nil Ratan Sircar Medical College and Hospital, Kolkata, West Bengal-700014, India

**Author notes:** Address of corresponding author: Dr. Nishant Chakravorty, Assistant Professor, School of Medical Science and Technology, Indian Institute of Technology, Kharagpur, Paschim Medinipur, West Bengal -721302, India, Contact no.

**Keywords:** β-hemoglobinopathies, Fetal hemoglobin, non-coding RNAs, Machine learning, lncRNA-miRNA-mRNA networks

## Abstract

Fetal hemoglobin (HbF) induction is considered to be a promising therapeutic strategy to ameliorate the clinical severity of β-hemoglobin disorders, and has gained a significant amount of attention in recent times. Despite the enormous efforts towards the pharmacological intervention of HbF reactivation, progress has been stymied due to limited understanding of γ-globin gene regulation. In this study, we intended to investigate the implications of lncRNA-associated competing endogenous RNA (**ceRNA**) interactions in HbF regulation. Probe repurposing strategies for extraction of lncRNA signatures and subsequent *in silico* analysis on publicly available datasets (**GSE13284, GSE71935** and **GSE7874**) enabled us to identify 46 differentially expressed lncRNAs (**DElncRNAs**). Further, an optimum set of 11 lncRNAs that could distinguish between high HbF and normal conditions were predicted from these DElncRNAs using supervised machine learning and a stepwise selection model. The candidate lncRNAs were then linked with differentially expressed miRNAs and mRNAs to identify lncRNA-miRNA-mRNA ceRNA networks. The network revealed that 2 lncRNAs (UCA1 and ZEB1-AS1) and 4 miRNAs (hsa-miR-19b-3p,hsa-miR-3646,hsa-miR-937 and hsa-miR-548j) sequentially mediate cross-talk among different signaling pathways which provide novel insights into the lncRNA-mediated regulatory mechanisms, and thus lay the foundation of future studies to identify lncRNA-mediated therapeutic targets for HbF reactivation.

## 1. Introduction

Fetal hemoglobin (α_2_γ_2_) is the predominant form of hemoglobin at the time of birth. The switching from γ-globin to β-globin gene expression occurs during ontogeny, leading to the replacement of fetal hemoglobin (HbF) to adult hemoglobin (HbA) in a gradual manner(Bank 2006). The complex globin gene switching mechanisms are fundamentally important to understand higher eukaryotic transcriptional regulation. Fetal hemoglobin represents less than 1% of total hemoglobin in adult individuals; and has low clinical relevance in normal physiology(Sankaran and Orkin 2013). However, congenital, acquired, and drug-induced strategies for HbF induction are reported to ameliorate the clinical manifestations of β-hemoglobin disorders such as Sickle cell anemia (SCA) and β-thalassemia disease by reducing the α-globin chain precipitation and providing an alternate form of functional hemoglobin. Therefore, understanding the intricate mechanisms of γ-globin gene regulation are expected tohave direct translational clinical relevance and hence lead to potential therapeutic advancements in β-hemoglobinopathies(Demirci, Leonard, and Tisdale 2020).

Numerous factors, covering both genetic and epigenetic alterations are known to influence the regulation of γ-globin genes in mature adults. Although certain hereditary conditions such as Hereditary persistence of fetal hemoglobin(HPFH), delta-beta thalassemia (δβ-thalassemia) and pathological conditions like Juvenile myelomonocytic leukemia(JMML) provide us with natural platforms to gain useful insights regarding γ-globin gene regulation in adulthood (Fluhr et al. 2017)(Wienert et al. 2015); however, a detailed understanding of the mechanisms to elevate γ-globin expression are in still primitive stage and attracts prime attention of the researchers throughout the world.

As the search for methods to elevate HbF levels to treat β-hemoglobin disorders such as sickle cell anemia (SCA), beta-thalassemia etc. continues, growing evidence suggests a major role of non-coding RNAs as modifiers of fetal hemoglobin expression(Sumera et al. 2020). Long non-coding RNAs (lncRNAs) -a major class of non-coding RNAs (ncRNAs) longer than 200 nucleotides, have become the focus of study in the recent years. Previously thought as transcriptional noise, these long non-coding RNAs are reported to exert their regulatory role at different levels including transcription, post-transcription and translation, and can modulate protein expression, thus playing pivotal roles in various physiological and pathological processes like cell division, cellular differentiation, embryonic development, carcinogenesis and neurodegenerative diseases (Kazemzadeh, Safaralizadeh, and Orang 2015). In the last decade, many studies have shown that lncRNAs have important implications in hematopoiesis and blood-related pathophysiologies (Xueqing Zhang, Weissman, and Newburger 2014)(Alvarez-Dominguez et al. 2014)(Nobili, Lionetti, and Neri 2016). Recently, it has reported that lncRNAs can interact with microRNAs (miRNA). miRNAs are major subsets of ncRNAs which belong to the category of small non-coding RNAs (sncRNAs) class(<200 nucleotides in length). They are known to regulate target gene expression by inhibiting mRNA translation or promoting mRNA degradation(Gebert and MacRae 2019). LncRNAs can function as competing endogenous networks (ceRNA) with mRNAs and small non-coding RNAs. They are reported to act as miRNA sponges that can sequester miRNAs and consequently inhibit their interactions with target mRNAs. LncRNAs can directly bind to the 3’UTR of mRNA, the binding site of miRNA, and thus preventing miRNA-mediated repression or degradation (López-Urrutia et al. 2019)(Paraskevopoulou and Hatzigeorgiou 2016). On the contrary, some miRNAs are reported to modulate lncRNA expression (Huang 2018). In addition, lncRNAs have been shown to act as precursors of the miRNAs and thus can regulate miRNA biogenesis in different pathophysiological conditions(Fernandes et al. 2019).

Recently, some studies have demonstrated the emerging role of lncRNAs in the regulation of fetal hemoglobin (HbF) expression(Ivaldi et al. 2018)(Jia et al. 2019). It is expected that identification of new lncRNAs would contribute substantially to our understanding of the regulatory mechanisms of the γ-globin genes and hence bring us closer to developing new therapeutic strategies to treat beta-hemoglobinopathies like β-thalassemia and SCA disease. Genome-wide association studies (GWAS) have found that three major gene loci – *HBB, HBSIL-MYB* intergenic region, and *BCL11A*, account for more than 20-45% of the HbF variations amongst various population (Uda et al. 2008). In the *HBS1L-MYB* locus, a 3bp deletion polymorphism in an enhancer sequence leads to the synthesis of a 1283bp transcript HMI-lncRNA. Upon knockdown of this lncRNA, gamma globin expression increased 200 folds. The binding site of this transcript and downstream mechanisms are yet to be elucidated but it has been hypothesized to interact with the *MYB* promoter to regulate HbF levels (Morrison et al. 2018). Another lncRNA encoded downstream of ^A^γ-globin, known as BGLT3, actively contributes to the increased expression of HbF. The precise mechanism for this action is by looping between the γ-globin gene and BGLT3 sequences and interaction with chromatin mediator complex (Ivaldi et al. 2018). These significant studies conducted till now are suggestive of the pivotal role played by long non-coding RNAs in facilitating the transcriptional control of γ-globin synthesis. However, to best of our knowledge no study has focused on the possible involvement of the lncRNA-miRNA-mRNA-signaling pathway network in HbF regulation and their effects on HbF induction so far.

In the present study, we’ve explored the possible molecular regulation of γ-globin expression by predicting intricate networks of lncRNA-miRNA-mRNA-signaling pathway axis involved in HbF regulation. We have obtained datasets GSE13284, GSE71935 and GSE7874 (Sankaran et al. 2008)(Helsmoortel et al. 2016)(Tanno et al. 2007) from the publicly available Gene Expression Omnibus (GEO) database and carried out *in silico* analysis to identify the differentially expressed lncRNAs. Supervised Machine Learning methods were employed to construct the lncRNA-based classifier to distinguish between high and normal HbF condition. Additionally, we performed a feature selection procedure to identify optimal set of lncRNAs that are possibly associated with high HbF level. We analyzed Gene Ontology (GO), Kyoto Encyclopedia of Genes and Genomes (KEGG) pathways and constructed the ceRNA network by connecting lncRNA-miRNA-mRNA-signaling pathway axis to gain new insights of intricate mechanisms of HbF regulation which could suggest novel targets of HbF reactivation in β-hemoglobinopathies. The overall workflow is schematically represented in Figure 1.

**Figure 1.**
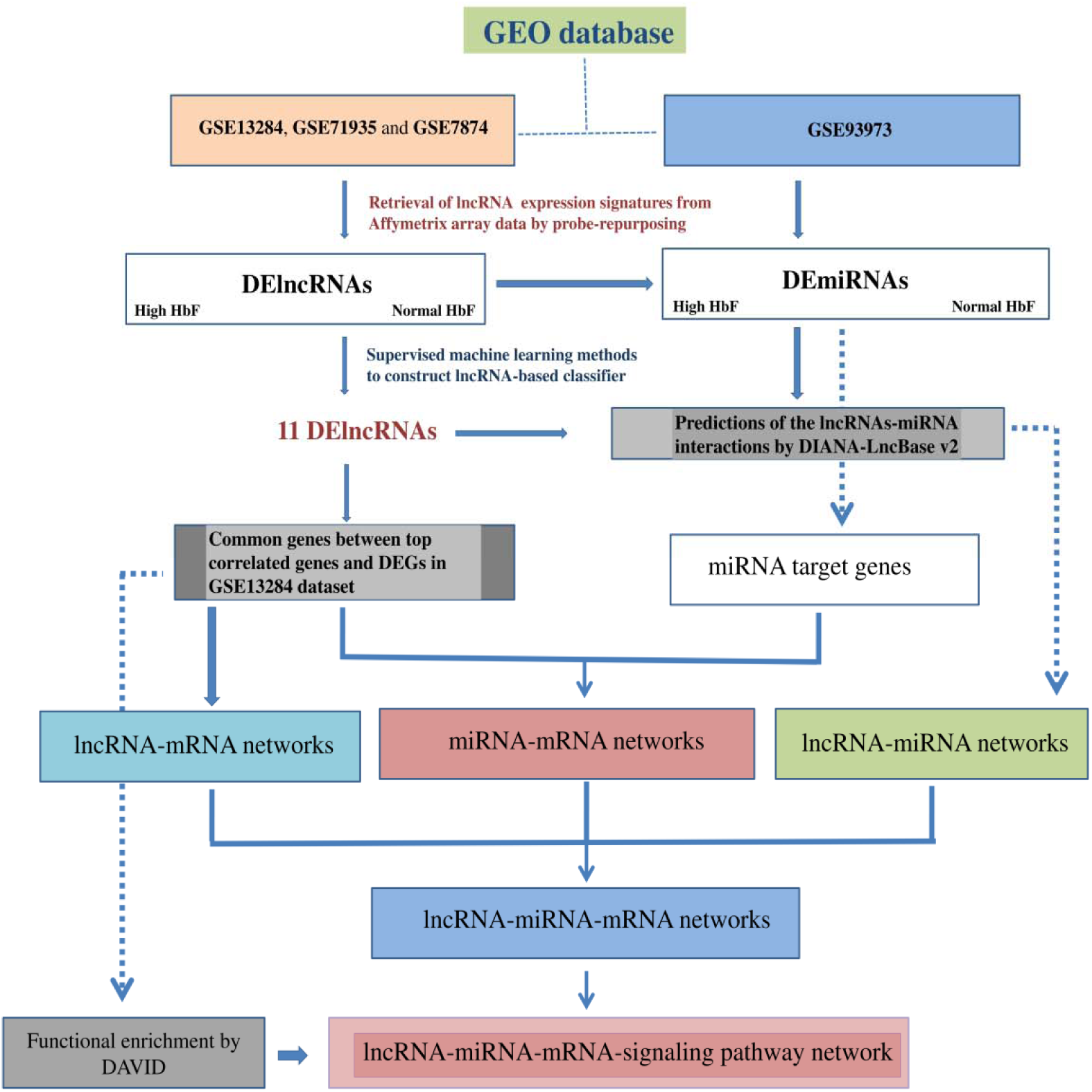
Flowchart depicting methodological workflow of predicted lncRNA-miRNA crosstalk in HbF regulation.

## 2. Results

### 2.1. LncRNA expression profiles on Affymetrix arrays

To extract lncRNA expression profiles, all datasets were transformed into probeID-centric data. A total of 54,675 entries were obtained for each dataset. By probe-repurposing and annotating the probe-sets (RefSeq and Ensemble annotations), we have obtained 2248 unique lncRNAs in our datasets (Supplementary file 1) which were utilized for further analysis. The probe sets with ambiguous annotations were excluded from our study.

### 2.2. Distinctive lncRNA expression between high HbF and normal HbF condition

The lncRNA expression patterns were compared between high HbF and normal HbF condition to identify the lncRNAs which are intricately associated with fetal hemoglobin regulation. Differentially expressed lncRNAs between two study groups were identified by performing Significant Analysis of Microarrays (SAM) analysis in training dataset, GSE13284. We obtained a total of 46 differentially expressed lncRNAs that had significant differential expression between the studied groups (Figure 2). Among the 46 deregulated lncRNAs, 20 lncRNAs were found to be up-regulated and the rest, 26 lncRNAs were identified to be down-regulated in patients (Supplementary file 2).

**Figure 2.**
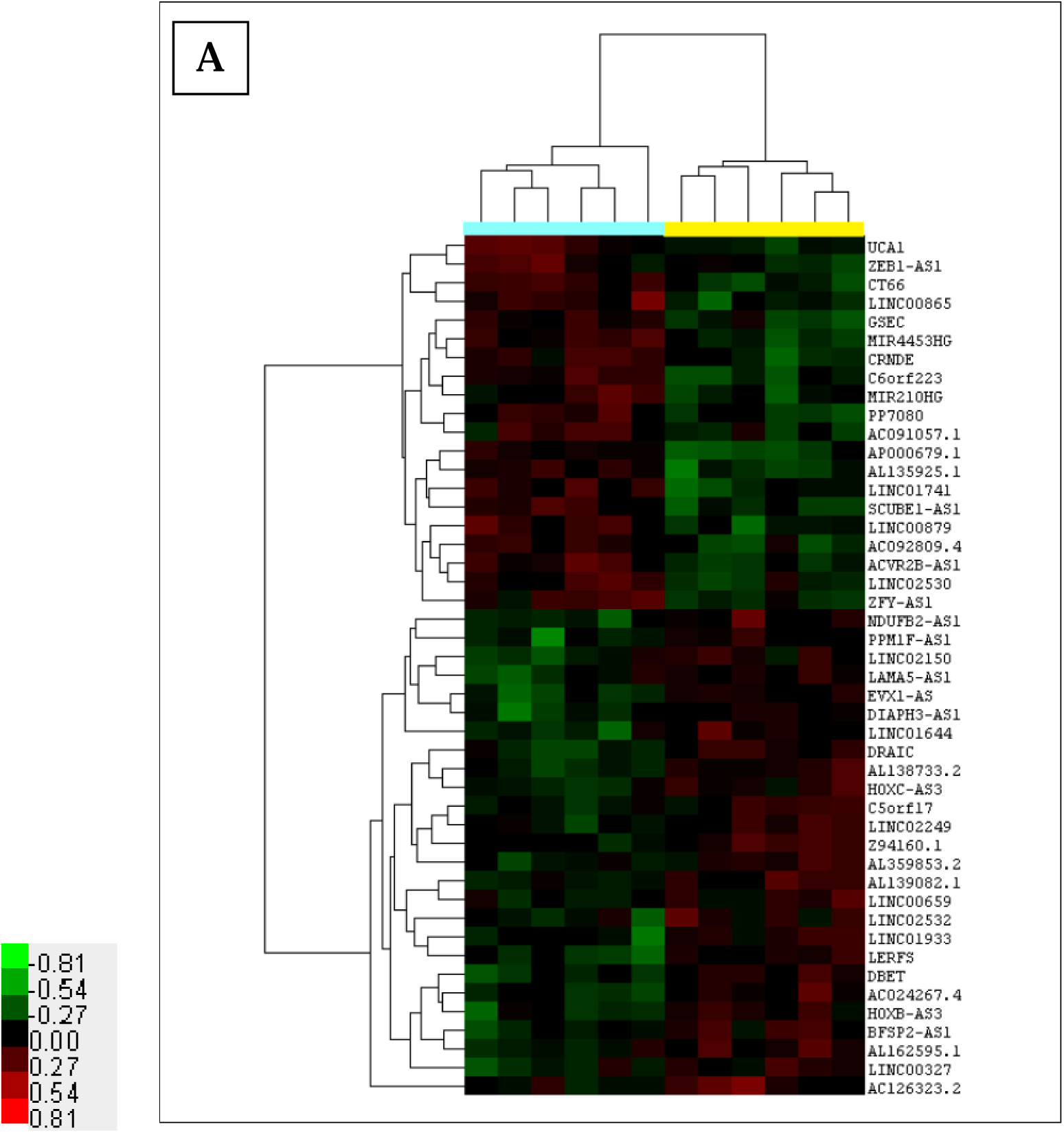
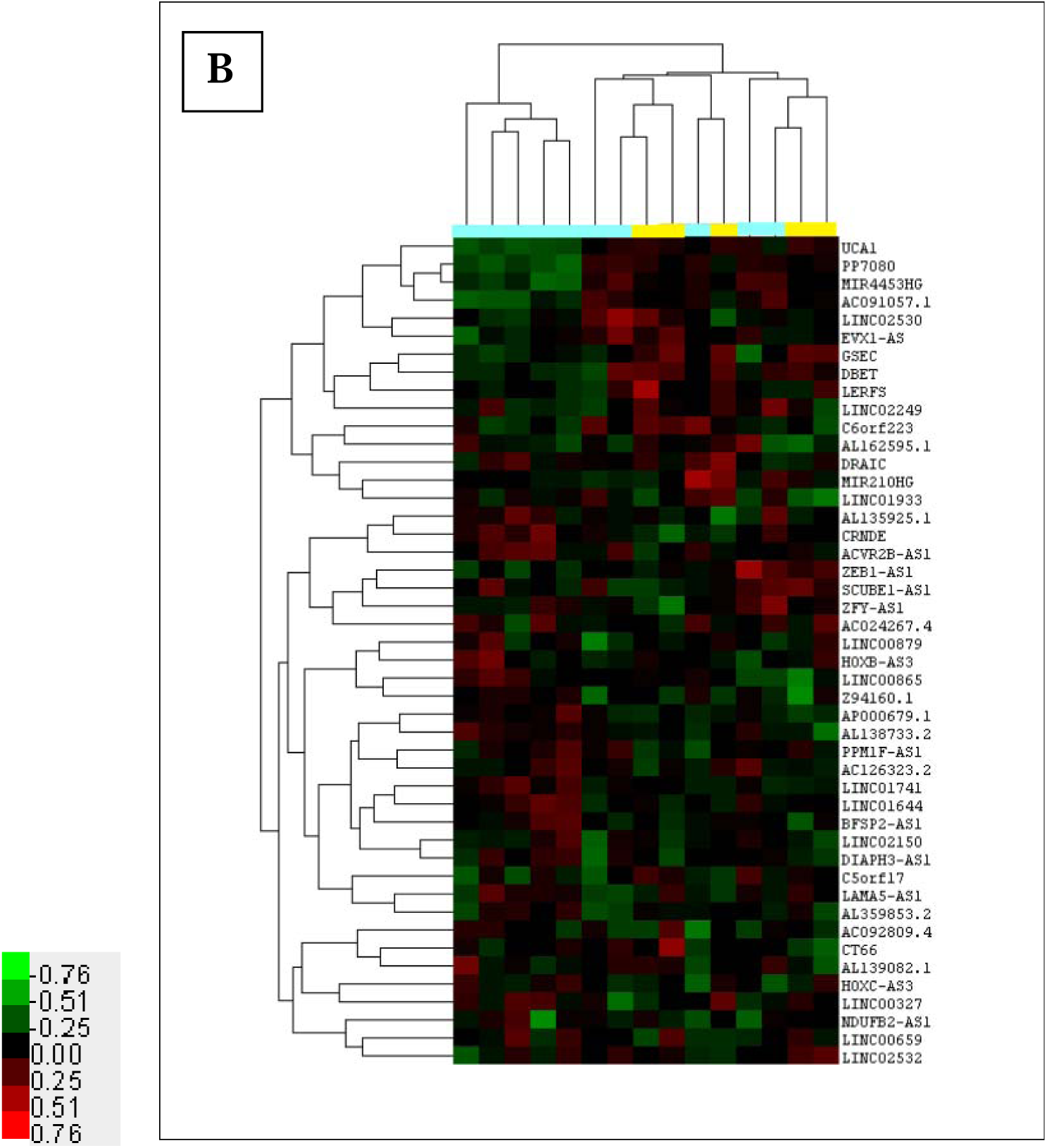
Differential expression of lncRNA probe sets between high HbF and normal condition. A total of 46 differentially expressed lncRNAs were identified to be significant in SAM analysis. **A**. The hierarchical clustering of lncRNA probe sets in training dataset, GSE13284. **B**. Validation of lncRNA probe sets in the discovery or test dataset, GSE7874.T-statistics with FDR<25% and Permutation of 1000 were performed to determine significant genes. Red= high expression; Green=low expression. Bar colors indicate sample type: yellow, high HbF; blue, normal HbF.

### 2.3. Supervised machine learning approach to determine optimal set of lncRNA, which could be involved in fetal hemoglobin regulation

In order to identify optimal combination of lncRNAs which could play a crucial role in fetal hemoglobin regulation, we performed an lncRNA-based classification and utilized feature selection methods with the46 differentially expressed lncRNAs using a supervised machine learning algorithm, SVM with radial basis function (RBF) kernel and stepwise selection model (Figure 3).

**Figure 3.**
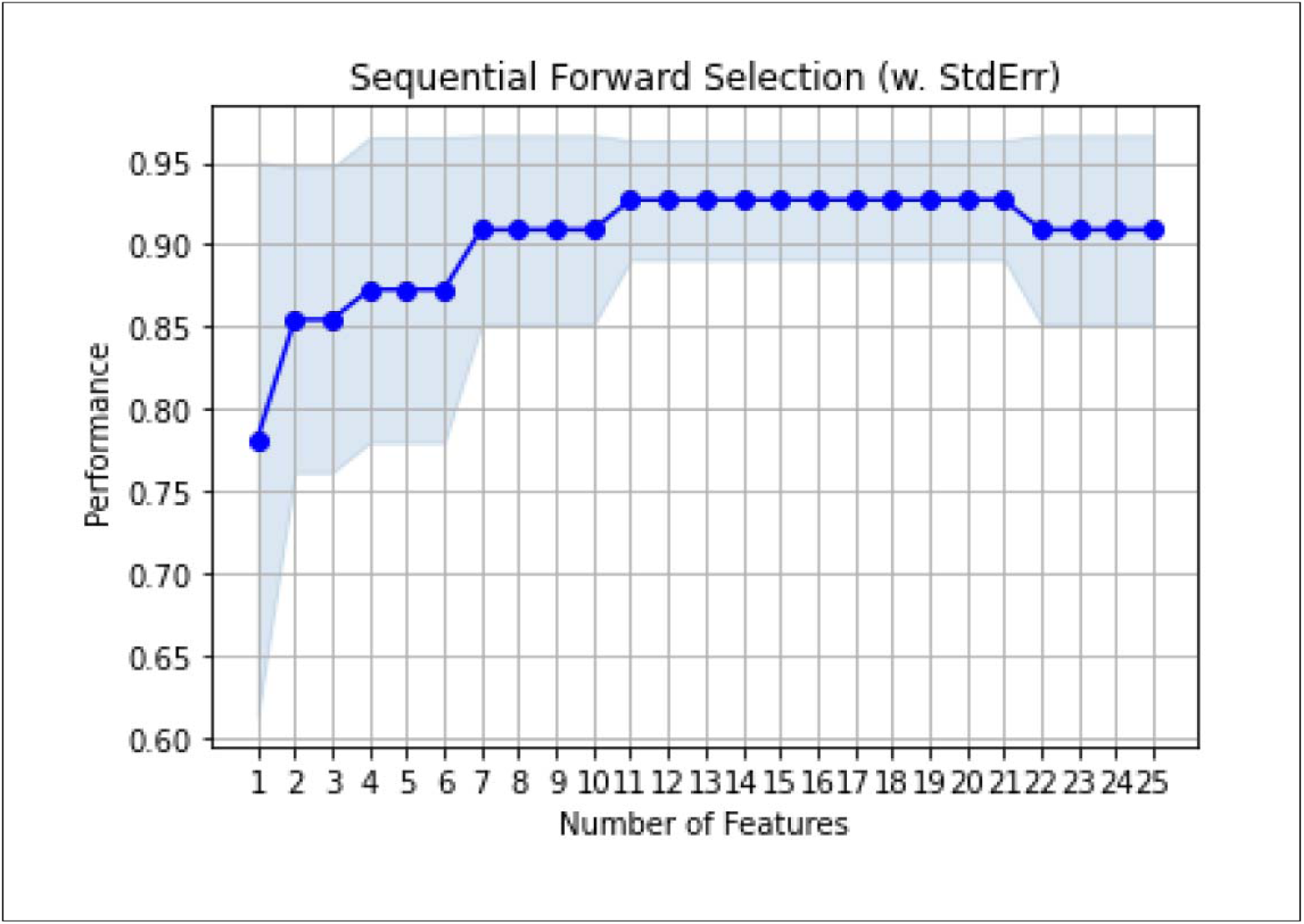
Showing features selection procedure to identify optimal set of lncRNA from differentially expressed lncRNAs. Stepwise selection method, involving forward selection was used.

A total of 11 (**C6orf223, AP000679.1, GSEC, UCA1, CT66, MIR4453HG, AL135925.1, ZEB1-AS1, AC091057.1, PPM1F-AS1** and **LINC02249**) out 46 differentially expressed lncRNAs were identified to be optimal combination which could form the lncRNA-based network for fetal hemoglobin regulation. Three machine learning classifiers, i.e. Random Forest, Support Vector Machine with linear kernel (SVM-L) and Support Vector Machine with RBF kernel (SVM-RBF), were tried, and we have calculated the performance metrics using the optimal set of 11 lncRNAs. The best results for the test set data were obtained for the SVM-RBF classifier with Accuracy and Precision of 89.47% and 93.33 %, respectively. The performance metrics for the test set data using all the classifiers is shown in Table 1.

**Table 1.** Performance matrix of different ML classifiers

| ML Classifiers | Accuracy | Precision | Recall | F1 score | AUC |
| --- | --- | --- | --- | --- | --- |
| Random Forest | 0.8421 | 0.8285 | 0.7948 | 0.808 | 0.8461 |
| SVM(Linear) | 0.7368 | 0.7023 | 0.7179 | 0.7076 | 0.7564 |
| SVM(RBF) | 0.8947 | 0.9333 | 0.8333 | 0.8642 | 0.8076 |

The Receiver Operating Characteristics (ROC) for all the classifiers employed in our study were plotted to show the discriminative power of each of this classifier. A comparison of the ROC curves computed for all the three classifiers is shown in Figure 4.

**Figure 4.**
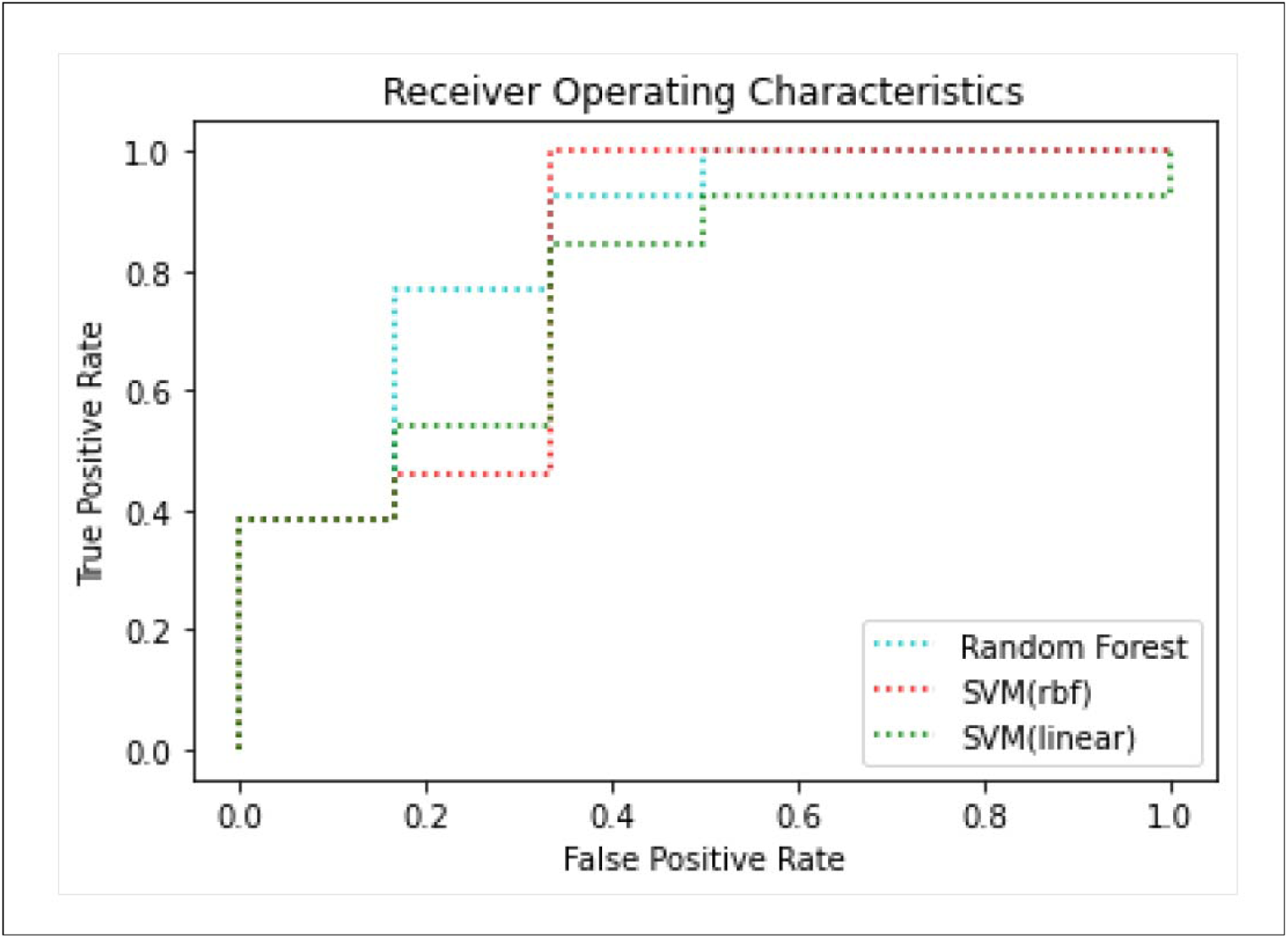
ROC curves of lncRNA-based classifiers in the discovery cohort.

### 2.4. Correlated differentially expressed genes which are important in lncRNA-mRNA networks construction

To investigate the functional implications of the optimal set of 11 lncRNAs, we performed a correlation test to identify the association between each of these 11 lncRNAs and mRNAs in GEO dataset, GSE13284. A list of 16,383 correlated genes was obtained for each lncRNAs. Of these co-expressed mRNAs, top (1%) positively and negatively correlated genes were selected for further study. In addition, 82 genes were found to be differentially expressed between high HbF and control group in the GSE13284 dataset (Sankaran et al. 2008) (Figure 5). The lists of differentially expressed genes are provided as Supplementary file 3.

**Figure 5.**
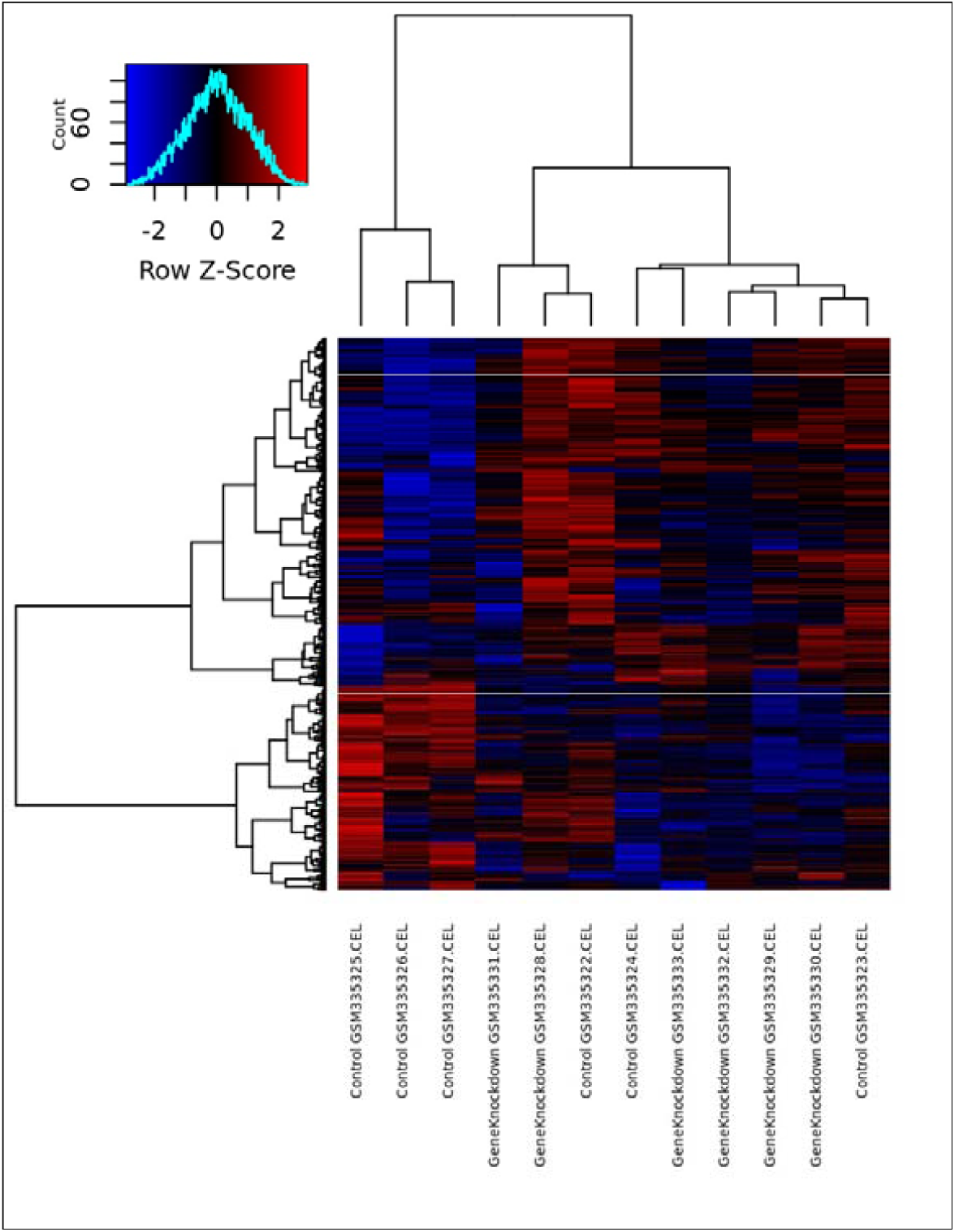
The Hierarchical clustering of differentially expressed mRNAs between high HbF and control group in GSE13284. Expression heatmap of differentially expressed genes was generated using R v3.6.3. “Red” and “Blue” colors represent above and below expression, respectively, relative expression.

Figure 5.The Hierarchical clustering of differentially expressed mRNAs between high HbF and control group in GSE13284. Expression heatmap of differentially expressed genes was generated using R v3.6.3. “Red” and “Blue” colors represent above and below expression, respectively, relative expression.

Common genes between top correlated genes and differentially expressed genes were used to construct lncRNA-mRNA networks.

### 2.5. Functional implications of optimal set of 11 lncRNAs

We performed GO term and functional enrichment analysis of shared mRNAs to gain insights regarding biological functions and cellular signaling pathways using DAVID resources. GO analysis revealed that the most enriched biological process (bp) was the glycolytic process (GO: 0006096), the most enriched cellular components (cc) was cytosol(GO: 0005829). The most enriched molecular function (mf) was protein binding (GO: 0005515) (Figure 6).

**Figure 6.**
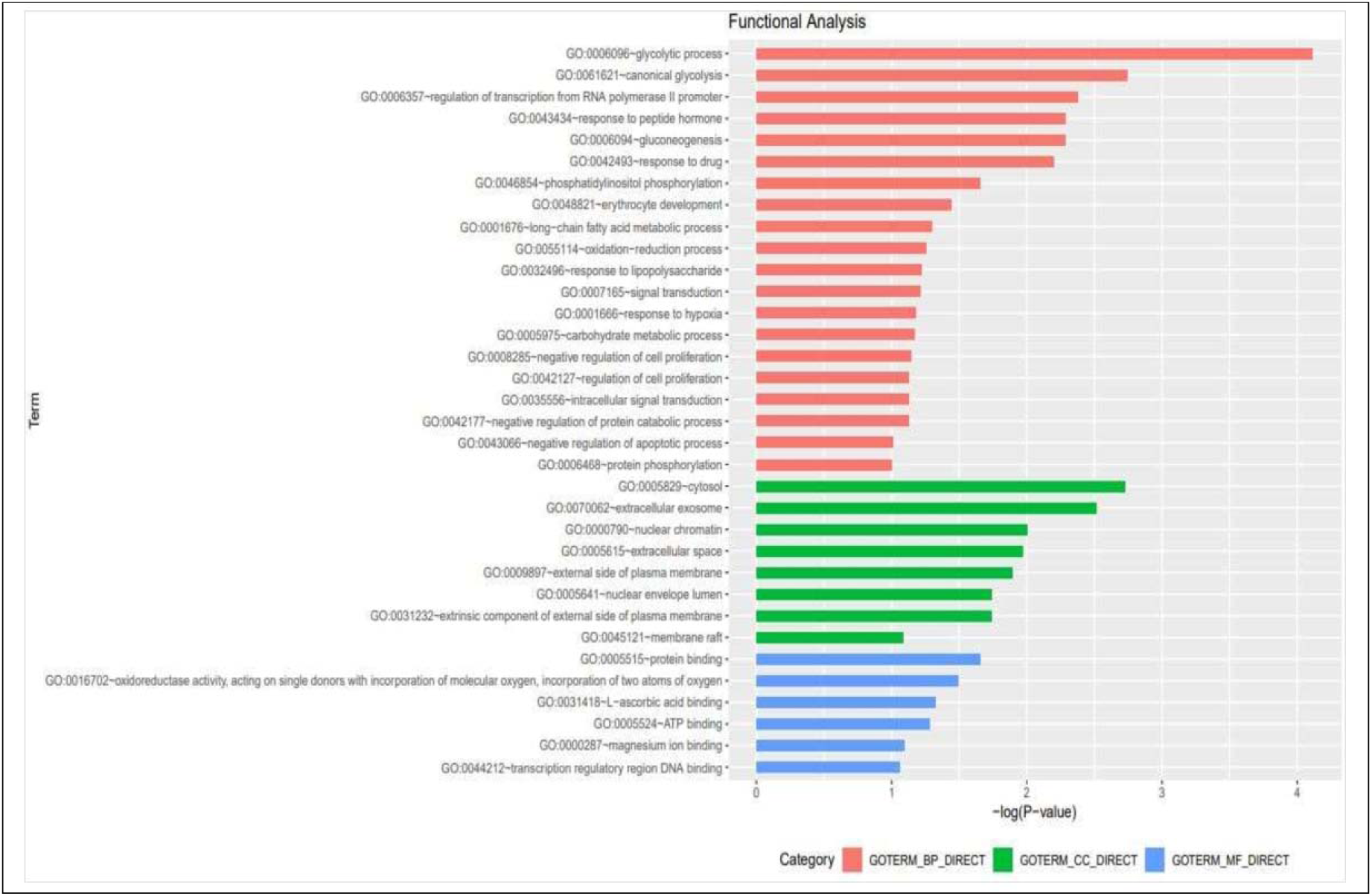
Gene Ontology (GO) annotation of differentially expressed correlated mRNAs, with a list top enriched genes (p<0.05), covering the domain of biological processes, cellular components and molecular functions. Enrichment values were –log10 (p-value) transformed.

Similarly, KEGG pathway analysis revealed 6 significant biological pathways that includes Glycolysis/Gluconeogenesis (hsa00010), Metabolic pathways (hsa01100), Biosynthesis of antibiotics (hsa01130), Biosynthesis of amino acids (hsa01230), Hematopoietic cell lineage (hsa04640) and Carbon metabolism (hsa01200) (Figure 7).

**Figure 7.**
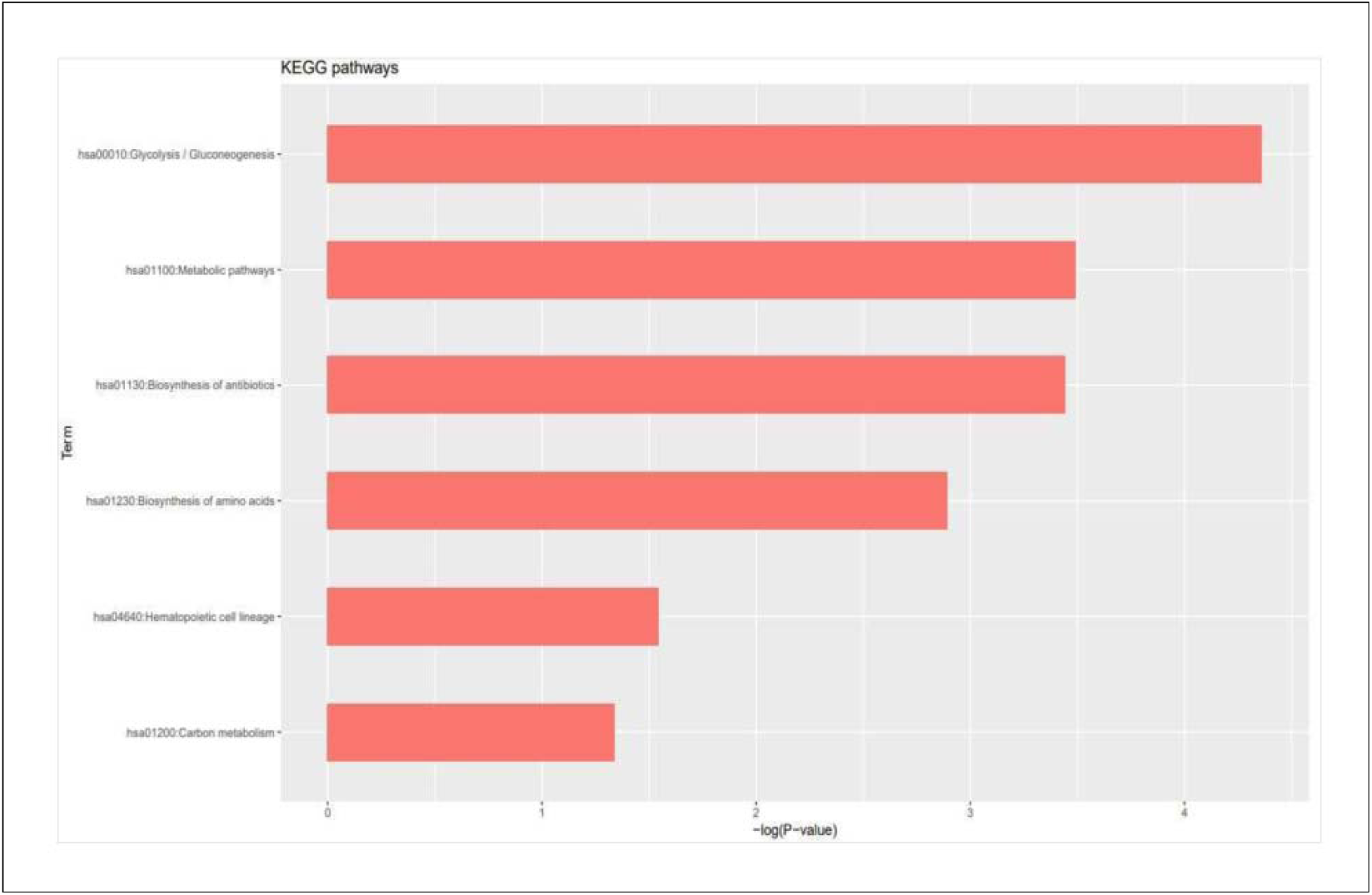
KEGG pathway analysis of common identified mRNAs. The 6 significant (p<0.05) signaling pathways are shown. All values are negative log transformed.

### 2.6. Construction of miRNA-mRNA regulatory networks

Based on differential miRNA expression analysis performed in GEO2R, 9 miRNAs (5 up-regulated and 4 down-regulated) were differentially expressed in high HbF vs. normal HbF conditions (Supplementary file 4). Out of these 9 differentially expressed miRNAs, 4 miRNAs (3 up-regulated and 1 down-regulated) had their targets common with list of differentially expressed common correlated mRNAs. A total of 14 genes and 4 miRNAs were found to have mutually opposite expression patterns, which were utilized for miRNA-mRNA networks construction (Figure 8).

**Figure 8.**
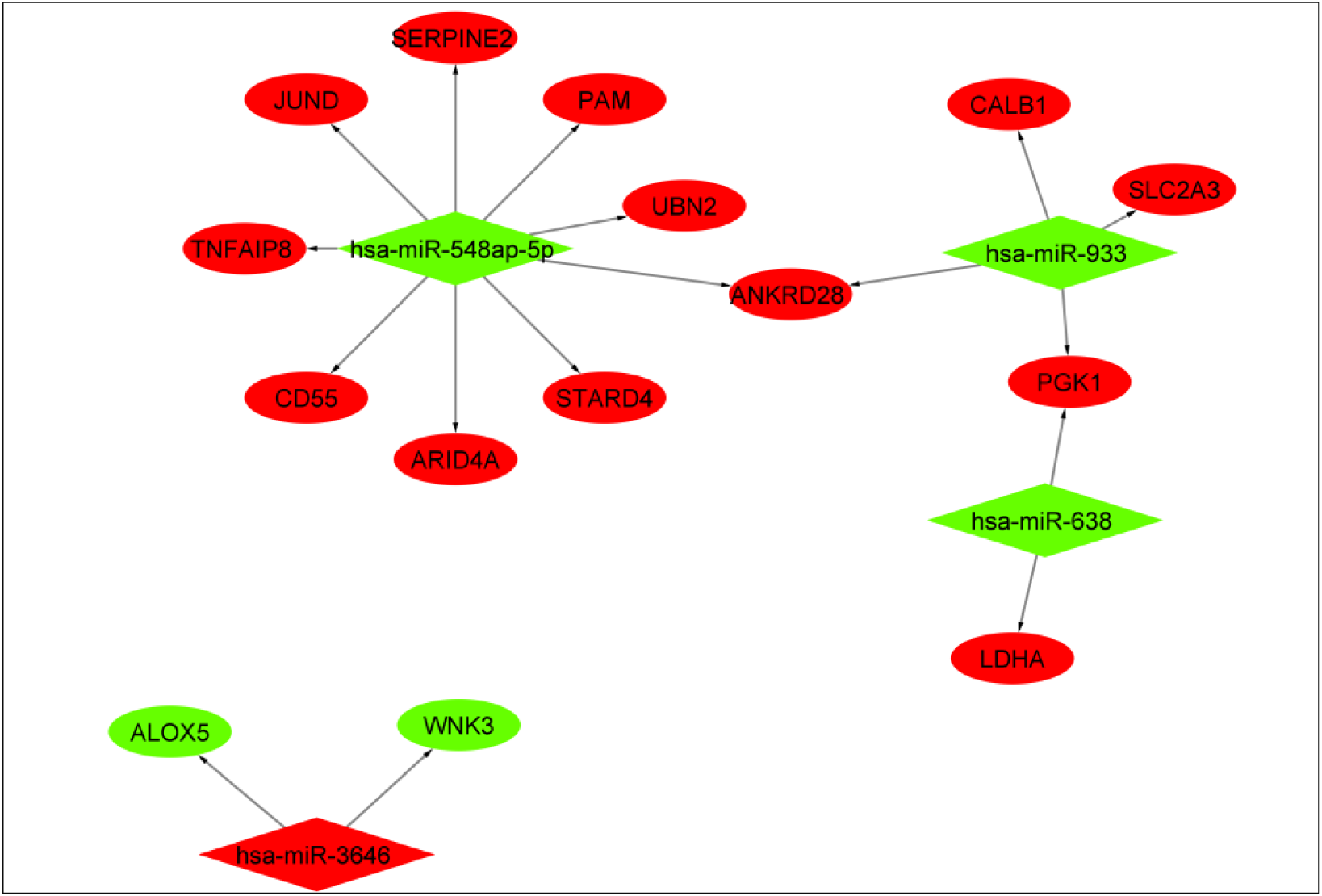
The figure is showing predicted miRNA-mRNA network. The network contains 19 nodes and 17 edges. The elliptical and diamond shapes represent mRNA and miRNA, respectively. Red and green colors indicate up-regulation and down-regulation, respectively. The directionality of the arrow is from the source to the target.

### 2.7. Prediction of lncRNA-miRNA networks

The interactions between identified lncRNAs and miRNAs were predicted using DIANA-lncBase v2. The differentially expressed miRNAs, which had shared targets with common correlated mRNAs and interacted with the identified optimal set lncRNAs, were selected for lncRNA-miRNA network construction (Supplementary file 5). Here, lncRNA-miRNA interactions with opposite expression are shown (Figure 9).

**Figure 9.**
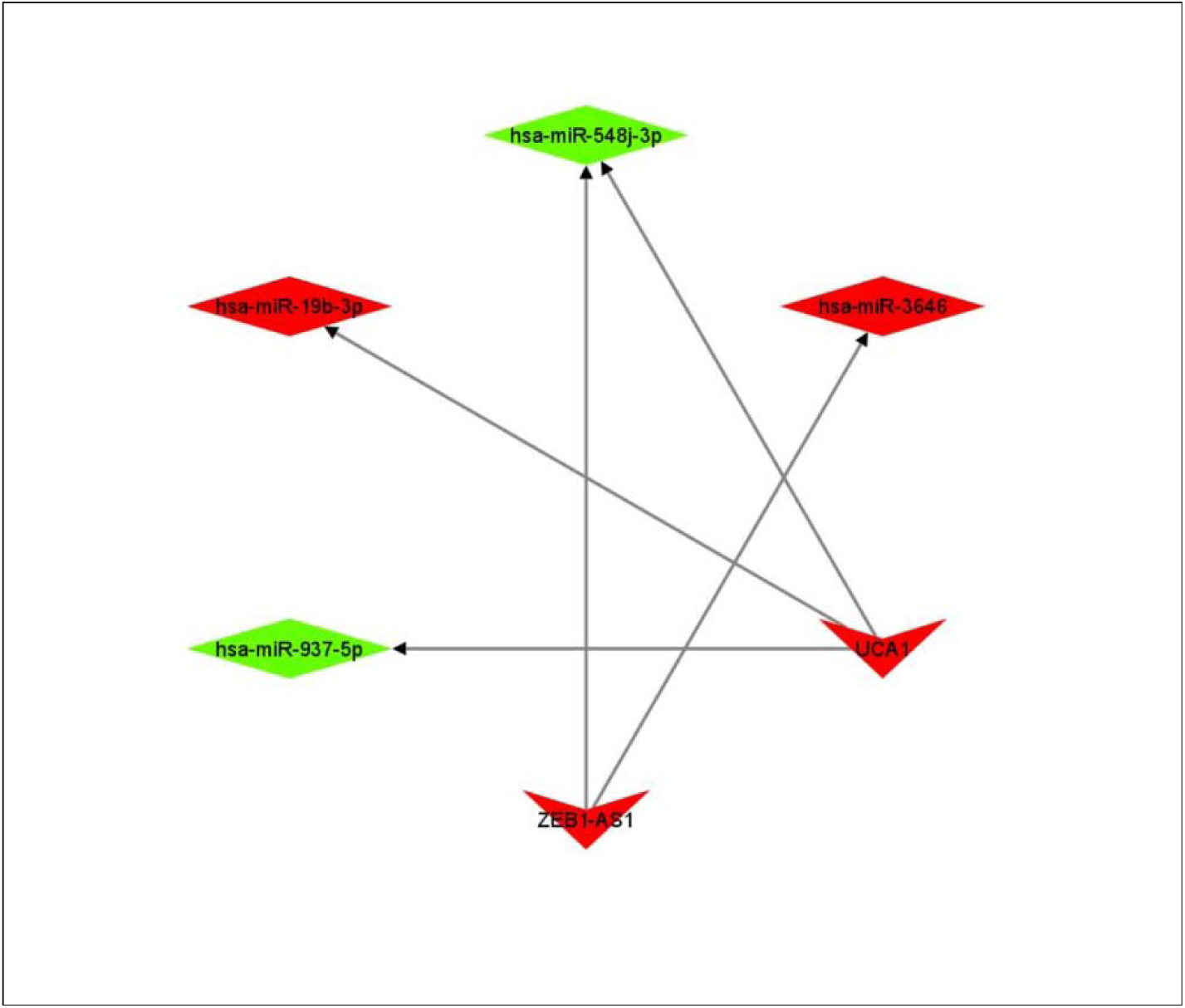
The figure is showing predicted lncRNA-miRNA network. The network consists of 6 nodes and 5 edges. The V-shapes and diamond represent lncRNA and miRNA, respectively. The red and green colors indicate up-regulation and down-regulation. Each arrow represents the interaction between lncRNA and miRNA, and the arrow is pointed from the source to the target.

### 2.8. Construction of lncRNA-miRNA-mRNA-signaling pathway network

Based on the predicted miRNA-mRNA, lncRNA-miRNA networks, we constructed lncRNA-miRNA-mRNA networks which were further linked with signaling pathways of the identified mRNA in the networks. In the figure 10, all the possible interactions among studied lncRNA. miRNAs and mRNAs are shown.

**Figure 10.**
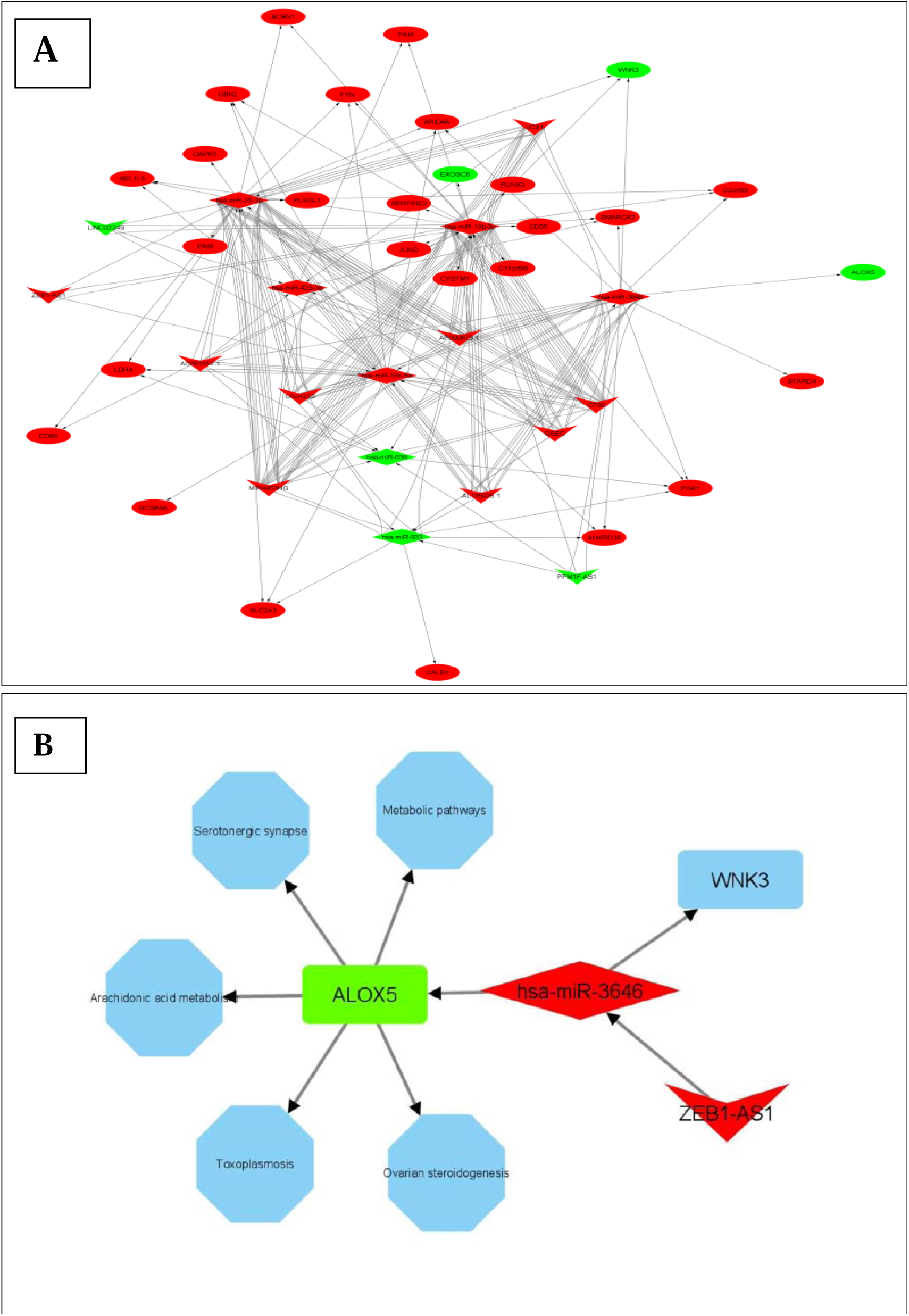
The interactions among lncRNA, miRNA and mRNA in high HbF vs. normal HbF condition. **A**. All possible networks among lncRNA, miRNA and mRNA in the studied groups. **B**. Predicted lncRNA-miRNA-mRNA-signaling pathway axis, which is important in HbF regulation. The V, Diamonds and elliptical shapes indicate lncRNA, miRNA and mRNA, respectively. Red= Up-regulation; Green= Down-regulation. The arrows show interactions and pointed from the source to the target.

Based upon the common link between lncRNA-miRNA and miRNA-mRNA networks, we have identified the lncRNA-miRNA-mRNA-signaling pathway axis which is predicted to be important for HbF up-regulation (Figure 10).

## 3. Discussion

Advancements in genomic and transcriptomic studies suggest towards the critical regulatory functions of non-coding RNAs in complex human diseases(Kazemzadeh, Safaralizadeh, and Orang 2015). Recent studies have highlighted the crucial roles of lncRNA in normal and malignant hematopoiesis (Ghafouri-Fard, Niazi, and Taheri 2021; Nobili, Lionetti, and Neri 2016). A study on the construction of an lncRNA-associated ceRNA network revealed two candidates, NR_001589 and uc002fcj.1, that are significantly expressed in individuals with high HbF levels. The study predicted that these lncRNAs promote the expression of HBG1/2 by sponging miRNA and induce HbF production (Jia et al. 2019). Microarray analysis in another study to profile lncRNAs, miRNAs and mRNAs in individuals having hereditary persistence of fetal hemoglobin has elucidated six lncRNAs that competitively sponge miR-486-3p, therefore elevating HbF level(Lai et al. 2017). Although few studies have performed the lncRNA, miRNA expression profiles, and attempted to provide preliminary evidences for the involvement of lncRNA-miRNA interactions in HPFH and β-thalassemia disease(Lai et al. 2017)(Fakhr-Eldeen, Toraih, and Fawzy 2019), the decisive role of lncRNA-miRNA-mRNA networks in HbF regulation remains largely obscure. Recent developments in high-throughput technologies and advancements in bioinformatics analysis, fueled by powerful Machine learning tools, have revolutionized the field of medical informatics. Here, we have explored microarray datasets involving high HbF vs. normal HbF condition, deposited in the GEO database, NCBI (Clough and Barrett 2016), and analyzed them using standard bioinformatics tools and machine learning algorithms to discern the complex regulatory networks involved in HbF induction. We have analyzed three independent microarray datasets (**GSE13284, GSE71935** and **GSE7874**) and identified 46 differentially expressed lncRNAs between high HbF and normal condition. To obtain optimal set of lncRNA that could distinguish between high HbF and normal HbF condition, we used different machine learning algorithms (Random Forest, Linear SVM and SVM-RBF), which further narrow down the numbers of lncRNA candidates. Comparative analysis revealed that the best results were obtained from the SVM-RBF model. A total of 46 differentially expressed lncRNAs were subjected to SVM-RBF and stepwise selection model, and we found a panel of 11 lncRNAs (**C6orf223, AP000679.1, GSEC, UCA1, CT66, MIR4453HG, AL135925.1, ZEB1-AS1, AC091057.1, PPM1F-AS1** and **LINC02249**) with accuracy of 0.8947 for distinguishing high HbF and normal control in the discovery cohort.

To identify the underlying molecular mechanisms and possible biological implications of these 11 candidate lncRNAs in HbF regulation, we attempted to correlate them with co-expressed mRNAs in GSE13284 dataset. We performed differential expression analysis (DEA) in the same dataset and correlated differentially expressed genes with 11 candidate lncRNAs. Functional enrichment analysis of the correlated genes revealed that these lncRNAs are involved in protein-binding (GO: 0005515), oxidoreductase activity (GO: 0016702), erythrocyte development (GO: 0046854), negative regulation of apoptotic process (GO: 0043066), etc. In an earlier study, Gabbianelli et al. 2008, have already reported that HbF reactivation inhibits cellular apoptosis in adult erythrocytes(Gabbianelli et al. 2008). Ma et al. demonstrated lncRNA HBBP1 increased γ-globin expression by activating ETS transcription factor ELK1 in HUDEP-2 cells (S.-P. Ma et al. 2021). A recent study has suggested the possible role of mRNA-binding protein in HbF reactivation in adult erythroid cells(Chambers et al. 2020). The fetal to adult hemoglobin switching is an important event in erythrocyte development(Sankaran, Xu, and Orkin 2010). KEGG pathway analysis revealed that major enriched pathways are involved in metabolic processes such as Glycolysis/Gluconeogenesis (hsa00010), Metabolic pathways (hsa01100), Biosynthesis of amino acids (hsa01230), Carbon metabolism (hsa01200) and Blood-cell development-related processes such as Hematopoietic cell lineage (hsa04640). Alterations in metabolic processes have already been reported in β-thalassemia patients with high HbF (Musharraf et al. 2017). In 2009, Testa et al. showed the role of stem cell factors in the reactivation of HbF in adults(Gabbianelli and Testa 2009).

Furthermore, we have also analyzed a separate miRNA expression dataset, **GSE93973**. Nine differentially expressed (adjacent p-value < 0.1, Log2FC > |0.8|) miRNAs (5 up-regulated and 4 down-regulated) were obtained, and their targets were predicted. Of these 9 deregulated miRNAs, 4 miRNAs shared the targets with common correlated differentially expressed mRNAs. Results revealed that hsa-miR-548ap-5p interacts with 9 out of 14 identified mRNAs in the network. Another miRNA, hsa-miR-933, is associated with 4 mRNAs in the network. Previous studies have demonstrated that intronic hsa-miR-933 plays a pivotal role in type II diabetes mellitus and neurodegenerative disease development (Bashar et al. 2020). To the best of our knowledge, no previous study has reported the possible link of this miRNAs to fetal hemoglobin development. Studies have identified that the loss of hsa-miR-638 facilitates cancer progression by targeting *SOX2*, an important transcriptional regulator of BCL11A (K. Ma et al. 2014)(Lazarus et al. 2018). In 2008, Sankaran et al. showed the role of BCL11A as a potential regulator of HbF expression (Sankaran et al. 2008). Another study investigated the effects of different deleterious SNPs on BCL11A and Fetal hemoglobin reactivation (Das and Chakravorty 2020). Furthermore, we observed hsa-miR-3646 targets two mRNA, WNK3 and ALOX5, in a separate network. Although, previous reports have suggested the role of hsa-miR-3646 in breast cancer progression (Tao et al. 2016); however no studies have been found which indicate the role of hsa-miR-3646 in HbF regulation. Our study suggests a possible link of this miRNA with HbF regulation.

Using DIANA-LncBase v2, we have predicted possible interactions between miRNAs and the 11 candidate lncRNAs identified. Finally, based upon the data from lncRNA-miRNA, miRNA-mRNA. lncRNA-mRNA networks, we have constructed the lncRNA-miRNA-mRNA regulatory axis of HbF expression, which was further linked with signaling pathways. The networks demonstrate the potential mechanism by which lncRNAs and miRNAs are involved in HbF regulation. As Fig 8 indicates, 2 lncRNAs (UCA1 and ZEB1-AS1) and 4 miRNAs (hsa-miR-19b-3p, hsa-miR-3646, hsa-miR-937 and hsa-miR-548j) have been identified as critical players in lncRNA-miRNA-mRNA networks. Recently, a study by Liu et al. has reported the role of lncRNA-UCA1 in heme biosynthesis and erythroid development (Liu et al. 2018). Another study reported that lncRNA-ZEB-AS1 promotes cell proliferation by regulating the Wnt signaling pathway, a crucial pathway in embryonic development(Lv et al. 2018). In a study, Mnika et al. demonstrated that hsa-miR-19b is associated with HU-induced HbF production in SCA patients(Mnika et al. 2019). All these reports are in accordance with our findings. Furthermore, by linking signaling pathways with the lncRNA-miRNA-mRNA networks, we have identified ZEB-AS1-hsa-miR-3646-ALOX5-signaling pathway axis, which might play an important role in HbF regulation and thus, could potentially contribute to therapeutic developments in β-hemoglobinopathies. Previous reports suggested the role of *ALOX5* in modulating the fetal inflammatory response in newborns (Costa and Castelo 2016). WNK signaling has also been reported to be associated with the deoxygenation of erythrocytes in normal physiology(Zheng et al. 2019). Infections pose a serious threat, and is reported to be the major cause of mortality and morbidity in thalassemic patients(Vento, Cainelli, and Cesario 2006). Although few studies have already reported the seroprevalence of *Toxoplasma* infection in β-hemoglobinopathies (Ferreira et al. 2017)(El-Tantawy, Darwish, and Eissa 2019), no study has reported the possible association of HbF regulation with toxoplasmosis. In our study, we have identified the possible link of ZEB-AS1-has-miR-3646-ALOX5 axis with toxoplasmosis. *ALOX5* gene has previously been reported to be involved in clearance of *Toxoplasma gondii* infection (Mashima and Okuyama 2015)(Aliberti, Serhan, and Sher 2002). Our study has linked all these signaling pathways, mRNA, miRNAs and lncRNAs, and suggested their possible role in HbF regulation.

In summary, this study represents a comprehensive analysis of possible molecular interactions among lncRNAs, miRNAs and mRNAs involved in HbF regulation. In this study, we have constructed an lncRNA-based classifier which could distinguish high HbF and normal condition with high specificity and sensitivity in discovery cohort. The candidate lncRNAs were then linked with differentially expressed miRNAs and mRNAs to predict lncRNA-miRNA-mRNA regulatory networks. Based on these networks, 2 lncRNAs (UCA1 and ZEB1-AS1) and 4 miRNAs (hsa-miR-19b-3p, hsa-miR-3646, hsa-miR-937 and hsa-miR-548j) identified to be crucial in HbF regulation. This analysis also revealed the ZEB1-AS1-hsa-miR3646-AlOX5-signaling pathways axis, which provides new insights about the molecular mechanisms of HbF regulation and suggests novel therapeutic targets to ameliorate disease severity in β-hemoglobinopathies.

## 4. Methods and materials

### 4.1. Data acquisition and processing

In the present study, three independent microarray datasets,**GSE13284** (Sankaran et al. 2008), **GSE71935** (Helsmoortel et al. 2016) and **GSE7874** (Tanno et al. 2007), which include high fetal hemoglobin and control group, were retrieved from the publicly available Gene Expression Omnibus (GEO) database. In addition, an independent miRNA expression dataset, **GSE93973** (Lai et al. 2017) concerning hereditary persistence of fetal hemoglobin (HPFH) and β-thalassemia minor with high HbF and control group was accessed from the NCBI GEO dataset for miRNA-mRNA network construction(Lai et al. 2017). For lncRNA-based classification, datasets were selected according to the following criteria: (1) dataset having high HbF vs. normal HbF expression profiles ;(2) dataset with sample no.>3; and (3) only Affymetrix HG-U133 plus 2.0 arrays expression profiles were included in our study. The raw (.CEL) files were downloaded, quantile normalized, Log2transformed and background corrected by using Robust Multichip Average (RMA, Windows Version)(Sahlabadi et al. 2018). Finally, we obtained probe ID-centric gene expression datasets. These datasets were further analyzed for lncRNA-based classification as described elsewhere(Xiaoqin Zhang et al. 2012).

### 4.2. Retrieval and analysis of lncRNA expression profiles

LncRNA expression profiles were obtained by probe repurposing of AffymetrixHG-U133 Plus 2.0 array based upon the NetAffx annotations following a standard lncRNA based classification pipeline(Xiaoqin Zhang et al. 2012).The annotations include gene name, gene symbol, Ensemble and RefSeq IDs for every probe set ID present in the dataset. LncRNAs extractions were based upon the Ensemble and RefSeq IDs of the probes. In case of RefSeq annotations, we extracted the probes labeled with ‘NR’ (NR stands for non-coding RNA in RefSeq database). Similarly, for probes with Ensemble gene IDs, we retrieved the probes labeled with ‘linc RNA’, ‘non-coding RNA’, ‘miscRNA’, etc. Following this, the other ncRNAs (miRNAs, snoRNAs, snRNAs, etc.) and pseudogenes were filtered out manually, retaining probes labeled with long non-coding RNAs only. Finally, 2248 unique lncRNAs corresponding to 4254 probe ids were obtained and used for downstream analysis.

### 4.3. Significance Analysis of Microarrays (SAM) analysis

The Significance Analysis of Microarrays (SAM) was performed using the R package(Chu et al. 2011)to evaluate the differentially expressed lncRNAs between the study groups (High HbF vs. Normal HbF). SAM analysis was carried out using stringent filters with a false discovery rate (FDR) <25% and permutations of 1000 to reduce ambiguity and increase accuracy. LncRNA probe sets with fold change (FC) > 1.5 and delta value 0.26 were defined as significantly different in SAM analysis. The differentially expressed lncRNAs were imported in Cluster view 3.0 (de Hoon et al. 2004) to carry out the Hierarchical Cluster Analysis (HCA).

### 4.4. LncRNA-based classification and prediction

Machine Learning Algorithms were utilized to build the lncRNA based classifier to distinguish between physiologically high HbF and normal HbF level in python 3.7 using scikit-learn package. All the three datasets, i.e. GSE13284, GSE7874 and GSE71935, were combined and implemented training-validation approach, in which 75% of the datawas employed for training and the rest 25% was used for testing. The list of lncRNAs obtained after Significance Analysis of Microarray (SAM) was fed to Sequential Forward Selector (SFS) using a supervised machine learning algorithm, i.e. Support Vector Machine (SVM) with radial basis function (RBF) kernel and 5 fold cross-validation to obtain the optimal combination of lncRNAs. Three machine learning classifiers, i.e. Random Forest, Support Vector Machine with linear kernel (SVM-L) and Support Vector Machine with RBF kernel (SVM-RBF), were used to determine the performance metricsusing the optimal set oflncRNAs. To determine the discriminative power of the lncRNA-based classifier, a receiver operating characteristics (ROC) curve was plotted, and the area under the curve (AUC) was calculated.

### 4.5. Construction of lncRNA-mRNA networks

To gain insights regarding the functional spectrum of lncRNAs, we performed a Pearson correlation test and calculated paired correlation coefficient between lncRNAs and mRNAs using GSE13284 dataset to construct the lncRNA-mRNA network. Differential expression analysis was performed forthe same dataset to obtain the differentially expressed genes between the studied groups. Correlated genes were then mapped against differentially expressed genes (DEGs), and common genes were extracted for further analysis.

### 4.6. Functional enrichment analysis

To evaluate the specific biological functions of these differentially expressed common correlated genes, we carried out Gene enrichment analysis using the Database for Annotation, Visualization and Integrated Discovery (DAVID) tool(Dennis et al. 2003). Additionally, enriched pathways associated with these genes were identified and visualized using the Kyoto Encyclopedia of Genes and Genomes (KEGG) pathway of DAVID software(Kanehisa and Goto 2000).

### 4.7. Construction of lncRNA-miRNA-mRNA regulatory networks

Differential expression analysis (DEA) was performed to obtain differentially expressed miRNAs (adjusted p-value < 0.1 and log2FC > |0.8|) between high HbF and normal conditions using publicly available miRNA expression dataset, GSE93973 (Lai et al. 2017), and the targets of these differentially expressed miRNAs were identified by using multiMiR R package and database(Ru et al. 2014). The multiMiR is a comprehensive collection of predicted and validated miRNA target prediction database which compiled data from 14 different databases. Differentially expressed miRNAs that have shared targets with common identified mRNA with opposite expression pattern were selected to construct miRNA-mRNA network. To obtain interactions among common lncRNAs and miRNAs, we used DIANA-LncBase v2(Karagkouni et al. 2020). The common differentially expressed miRNAs were mapped against the optimal set of selected 11 lncRNAs for miRNA-lncRNA network constructions. Finally, based on miRNA-mRNA. lncRNA-miRNA and miRNA-mRNA-signaling pathway networks, we constructed lncRNA-miRNA-mRNA-signaling pathway network. All networks were visualized using Cytoscape v3.1(Shannon et al. 2003).

## Acknowledgements

Authors would like to thank Indian Institute of Technology Kharagpur for providing the infrastructural support. Mr. Motiur Rahaman would like to thank the Ministry of Education for financial support.

